# Characterization for Drought Tolerance and Physiological Efficiency in Novel Cytoplasmic Male Sterile Sources of Sunflower (*Helianthus annuus* L.)

**DOI:** 10.1101/400531

**Authors:** Vikrant Tyagi, Gurpreet Kaur, Prashant Kaushik, Satwinder Kaur Dhillon

**Affiliations:** Department of Plant Breeding and Genetics, Punjab Agricultural University, Ludhiana 141004, India; Instituto de Conservación y Mejora de la Agrodiversidad Valenciana, Universitat Politècnica de València, Valencia 46022, Spain

**Keywords:** sunflower, cytoplasmic male sterility, combining ability, drought, water stress

## Abstract

Sunflower is sensitive to drought and its hybrids have a limited cytoplasmic diversity. The wild cytoplasmic sources of sunflower are not well exploited to their potential for drought tolerance and hybrid development. In this respect, we carried out a Line × Tester based genetic study using 19 sunflower genotypes representing, 13 cytoplasmic male sterile (CMS) lines from wild and conventional sources, 2 maintainer lines, and 4 restorer lines. The CMS and maintainer lines were crossed with restorer lines to develop sixty one-way F1 hybrids. The parents and their hybrids were evaluated under two water regimes viz., normal irrigated and water stress. A total of twelve important plant descriptors were studied over a period of two years. The significant differences were observed between parents and hybrids in both water regimes. Hybrids were higher in average values for all the descriptors than parents. The role of female parent was more prominent in the expression of traits in hybrids as compared to male parents. The CMS sources varied significantly regarding seed yield per plant and other physiological traits. Proline content was three times higher in parents and their hybrids under water stress, and it was not correlated with any other descriptor. Accession CMS-PKU-2A was identified as the best general combiner for leaf area and specific leaf weight. Whereas, CMS-234A was the best general combiner for biological yield and photosynthetic efficiency under both the conditions. The cross combinations CMS-ARG-2A × RCR-8297, CMS-234A × P124R, and CMS-38A × P124R were found significant for biological yield, seed yield and oil content under both environments. Overall, this study provides useful information about the cytoplasmic effects on important sunflower traits and drought stress tolerance when used in the different combinations.

## 1. Introduction

Sunflower (*Helianthus annuus* L.) is a commercially important oilseed crop, and its oil is comparable to virgin oil olive for health promoting benefits [1,2]. Botanically, sunflower is a cross-pollinated and self-incompatible plant [3]. Moreover, it out crosses freely with its wild relatives [4]. During the past few decades, the area and production of sunflower have increased because of its day length neutrality, a wider adaptability, and responsiveness to added inputs [5]. Similarly, these attributes of sunflower promote sustainable crop production system when sunflower is used with other crops in crop rotation [6]. The central component of sunflower breeding is the development of hybrids through cytoplasmic male sterility (CMS) system [7]. CMS is the absence of fertile pollen in the plant hence eliminates the need for manual emasculation for the development of a successful cross combination. CMS based hybrids system has been extensively distributed throughout the plant kingdom and well exploited for achieving yield targets, disease resistance, and drought tolerance [8-10].

CMS system in sunflower was initially discovered by in the 1970s [11] and later in the successive year, its fertility restoration genotype was also identified [12]. The hybrid system based on a CMS commonly comprises of a three-line system: the male sterile genotype (A line), which is maintained by maintainer (B line) usually with a fertile cytoplasm, and a restorer (R line) for restoring the male fertility in hybrids via dominant fertility restorer genes [8,13]. Earlier efforts of sunflower breeders using this system have resulted in the significant increase of the production and quality [14,15]. Drought is among the biggest crop production challenges of the 21st century and sunflower is not an exception to drought stress [16]. The yield losses due to drought are prominent in sunflower[17,18]. Especially, the exposure during anthesis and dough stages can result in up to 80% crop losses[19]. Wild relatives of cultivated sunflower are among the potential source of important genes for the drought tolerance[20]. The genus Helianthus has sufficient diversity, with around 51 species and 19 subspecies fitting to the genus[21]. The characterization of the genotypes based on relative water content, leaf water potential, photosynthetic efficiency, and proline content are pivotal to any water stress screening experiment [22,23]. Similarly, proline which is an important plant osmolyte and its accumulation in the leaves during prolonged drought spells is considered to provide drought tolerance [24].

In modern sunflower hybrids, PET-1 is the most common source of cytoplasm genome[7]. This homogeny in the cytoplasm of most of the modern hybrids might lead to colossal yield losses as happened to other crops e.g. Maize [25,26]. To reduce such kind of disaster in sunflower the necessity of diversification in the CMS source is inevitable. Consequently, to diversify the cytoplasmic base, attempts have been made and several new cytoplasmic sources have been identified [27]. Further, in some instances, the negative effects of cytoplasmic and nuclear interaction resulted in the reduced chlorophyll content and photosynthetic efficiency [28]. In contrast, a positive effect of cytoplasmic nuclear interaction was also reported on oil content [29]. Therefore, the influence of cytoplasmic effects on important agronomic traits needs to be understood more precisely.

Previously, using only molecular diversity based criterion a study consisting of 28 CMS lines was as carried out [30]. Hitherto, an extensive study on drought tolerance, trait performance, and correlation among traits is missing. Therefore, in our study, we have used the 13 CMS (A) lines and 2 maintainer lines (B). Further, these A and B lines were crossed with 4 restorer lines (R) to develop 60 one-way F_1_ hybrids. These 60 hybrids were produced with a Line x Tester matting design to obtain a detailed information regarding general and specific combiners along with gene actions involved [31,32]. The parents and their hybrids were evaluated under normally irrigated and water stress setting for two consecutive years.

## 2. Results

### 2.1 Variation in Parents and their Hybrids

In the UPGMA analysis, more similar ones are grouped together in the same cluster [33]. The cluster analysis showed that the alloplasmic line CMS-XA was distinct from the rest of the accessions (Figure 1). While the six allopasmic lines were clustered together (Figure 1). Moreover, out of four, three testers were clustered in the same cluster (Figure 1).

**Figure 1.**
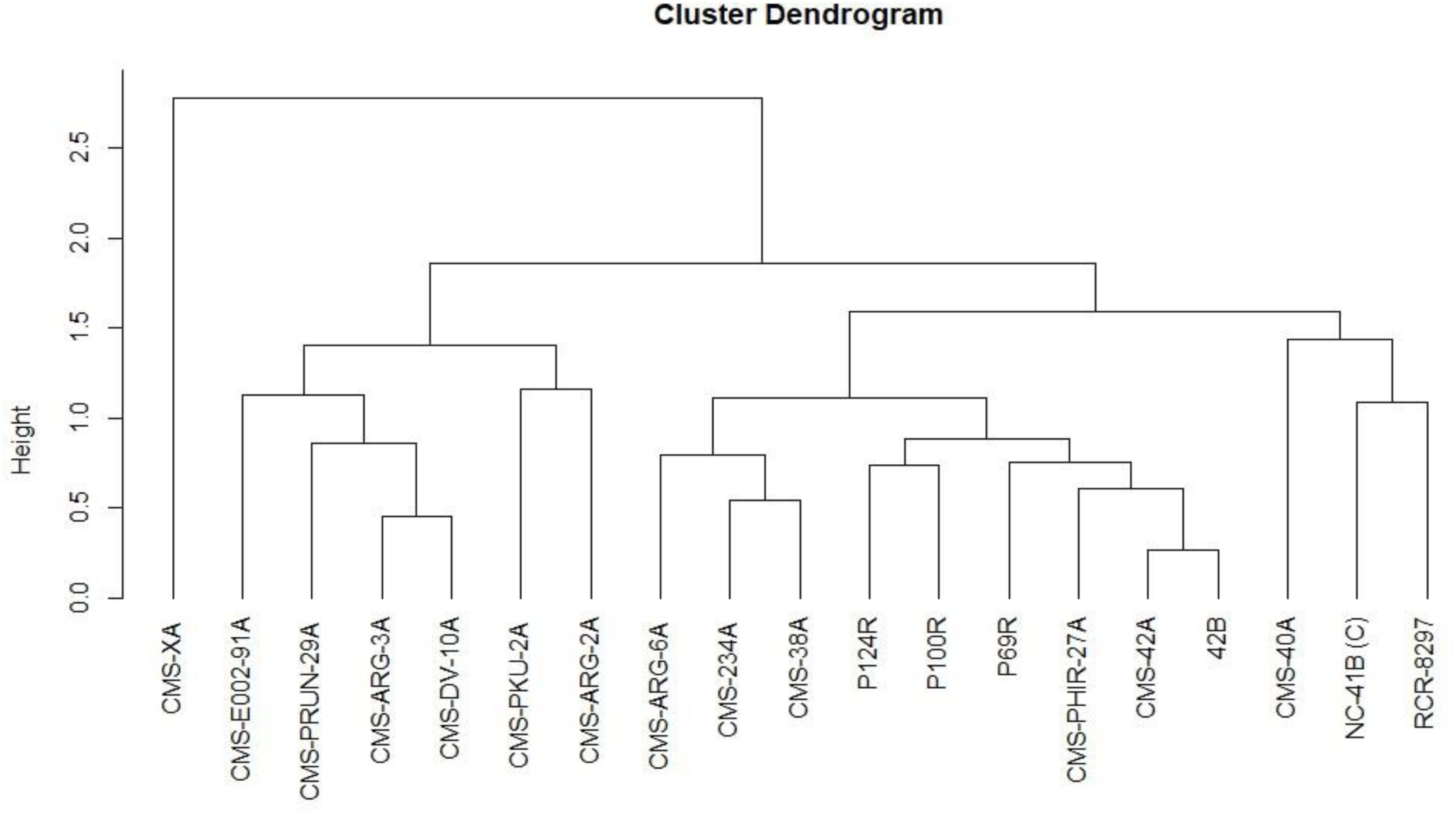
UPGMA clustering of parental genotypes into groups based on log-normalized descriptors values. The cophenetic correlation coefficient of clustering is 0.8.

There were highly significant differences (*P* < 0.05) for the mean values of groups of parents and hybrids (normal and water stress environment), for all of the 12 descriptors studied (Table 3). Parent and their hybrid combinations performed better under normal conditions than in the stress environment (Table 1). Likewise, hybrids were with more leaves, higher harvest index, seed yield, and oil content. Whereas, under water stress, parents and hybrids overlapped for harvest index and proline content (Table 1). The proline content was almost three times more in water-limited filed. Whereas, the photosynthetic efficiency remained to be unaltered under normal and drought condition (Table 1).

**Table 1.** Variation of parameters under normal and water stress environment.

| Descriptors | Parent (n=19) | Parent Stress | Hybrid (n=60) | Hybrid Stress | F-ratio | Probability |
| --- | --- | --- | --- | --- | --- | --- |
|  | Mean<sup>a</sup> | Mean<sup>a</sup> | Mean<sup>a</sup> | Mean<sup>a</sup> | F-ratio | Probability |
|  | (range) | (range) | (range) | (range) |  |  |
| Number of leaves | 22.31 b<br>(20.55-24.05) | 19.50 a<br>(17.75-21.25) | 24.65 b<br>(23.66-25.64) | 24.10 b<br>(23.11-25.08) | 9.61 | >0.001 |
| Leaf area (m<sup>2</sup>) | 0.62 b<br>(0.52-0.72) | 0.57 ab<br>(0.47-0.67) | 0.53 a<br>(0.47-0.58) | 0.45 ab<br>(0.39-0.50) | 3.98 | 0.0092 |
| Specific leaf weight (g) | 1.48 c<br>(1.25-1.71) | 1.38 bc<br>(1.16-1.62) | 1.15 ab<br>(1.02-1.28) | 1.00 a<br>(0.87-1.13) | 5.86 | 0.008 |
| Leaf area index | 3.41 b<br>(2.88-3.95) | 2.98 ab<br>(2.45-3.15) | 2.92 ab<br>(2.62-3.22) | 2.45 a<br>(2.15-2.75) | 3.82 | 0.0113 |
| Leaf water potential (MPa) | -2.38 b<br>(-2.32-(-2.15)) | -2.83 a<br>(-3.01-(-2.84)) | -2.23 b<br>(-2.53-(-2.23)) | -2.93 a<br>(-2.98-(-2.68)) | 48.7 | >0.001 |
| Relative leaf water content (%) | 68.52 c<br>(64.40-72.65) | 55.10 a<br>(50.97-59.23) | 65.97 c<br>(63.65-68.30) | 60.90 b<br>(58.56-63.21) | 10.29 | >0.001 |
| Photosynthetic efficiency | 36.60 a<br>(34.92-38.25) | 36.41 a<br>(34.74-38.08) | 35.58 a<br>(34.65-36.52) | 34.36 a<br>(33.43-35.30) | 2.76 | 0.0439 |
| Proline content (mg/g of dw) | 0.47 a<br>(0.36-0.57) | 1.58 b<br>(1.48-1.68) | 0.46 a<br>(0.40-0.52) | 1.51 b<br>(1.45-1.57) | 284.68 | >0.001 |
| Biological yield | 310.05b<br>(82.01-628.33) | 204.57a<br>(52.17-522.33) | 329.97b<br>(139.17-758.33) | 323.65b<br>(136.70-678.02) | 38.67 | >0.001 |
| Harvest index | 10.68 a<br>(7.80-13.57) | 17.81 b<br>(14.93-20.70) | 19.66 b<br>(18.04-21.28) | 17.97 b<br>(18.03-19.60) | 9.63 | >0.001 |
| Seed yield/Plant (g) | 25.88 a<br>(22.05-29.72) | 25.07 a<br>(21.23-28.90) | 52.90 c<br>(50.74-55.06) | 40.81 b<br>(38.66-42.97) | 81.83 | >0.001 |
| Oil content (%) | 29.38 b<br>(28.22-30.54) | 27.30 a<br>(26.13-28.45) | 31.23 c<br>(30.57-31.88) | 30.61 bc<br>(29.96-31.26) | 12.47 | >0.001 |

<sup>a</sup>The alphabet (a,b,c etc.) show significant differences based on the Student-Newman-Keuls test

The analysis of variance for combining abilities of the twelve descriptors studied in a line x tester (15 x 4) design is presented in Table 2. The mean squares due to treatments were highly significant for all the traits except for the proline content (Table 2). Likewise, the mean squares due to lines (female), testers (males), and female x male interactions were recorded highly significant for all the traits under both the environments and also, over the years (Table 2). Parentals and their hybrids had a highly significant (P ≤ 0.01) general combining ability (GCA) and specific combining ability (SCA) effects, for all the traits and under both the environments (Table 2). The ratios of GCA/SCA effect were >0.5 for biological yield and oil content under normal water environment, while under water stress environment leaf water potential, photosynthetic efficiency and harvest index, suggesting the predominance of additive over non-additive genetic effects. This ratio was <0.5 for most of the traits studied in both environments, implying a significant role of non-additive genetic effect on these traits (Table 2).

**Table 2.**
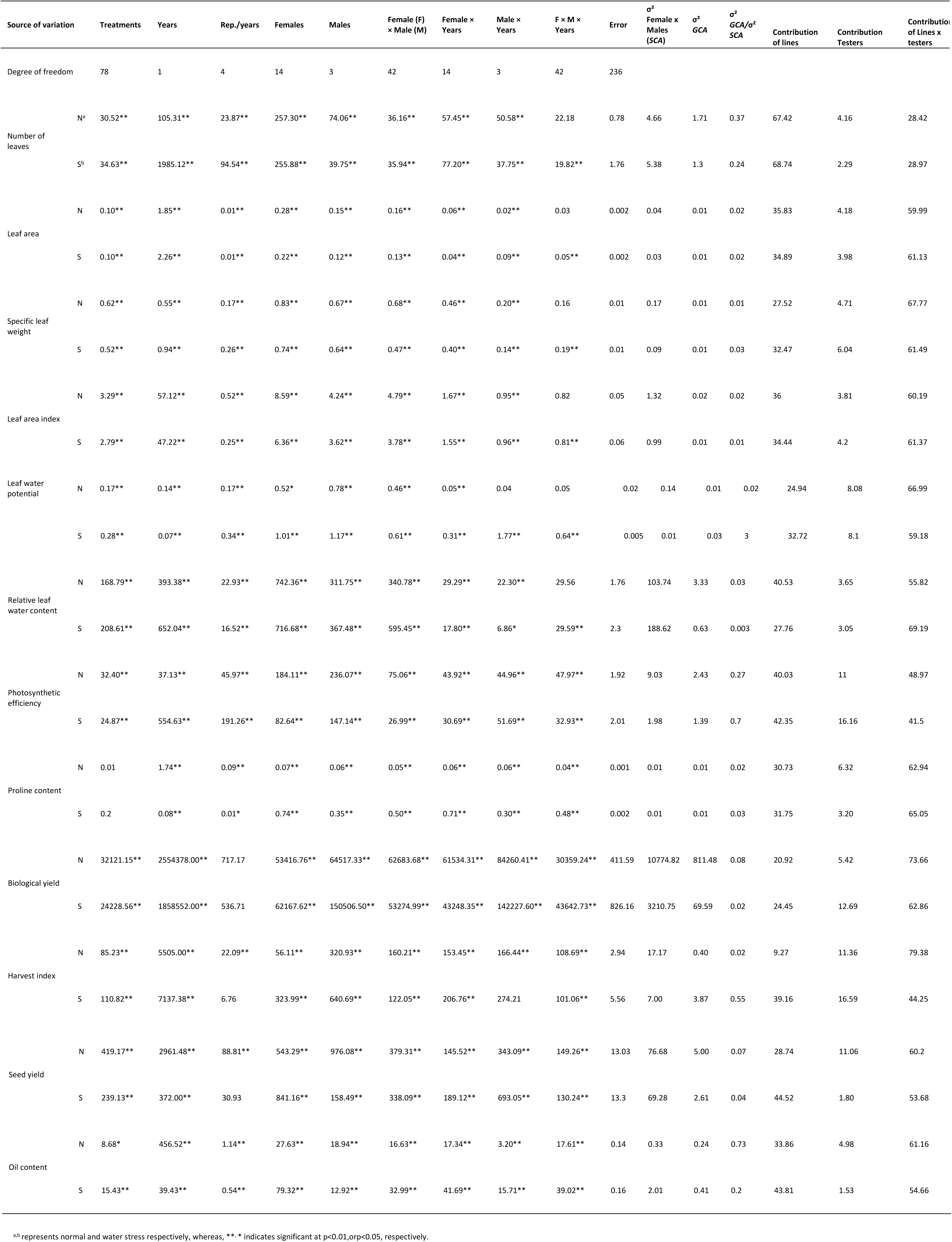
Analysis of variance for combining abilities (GCA and SCA) for twelve traits in sunflower across two environments.

The proportional contribution of parent and their interactions, the contribution of female parents were observed to be higher compared to testers (males) irrespective of the environments (Table 2). However, their overall interaction component (lines x testers) had a higher proportional contribution for traits *viz.,* leaf area, specific leaf weight, leaf area index, leaf water potential, relative leaf water content, proline content, seed yield, harvest index and oil content (Table 2).

### 2.2. Combining Ability Estimates

The estimates for the general combining ability (GCA) of the parental genotypes (19) under normal and under water stress environment for all 12 plant descriptors is provided in Table 3. The detailed estimates of specific combining ability (SCA) effects of hybrids for the all the traits under two different water regimes are presented in Table S1.

**Table 3.** The estimates for the general combining ability (GCA) of the parental genotypes (19) under normal and under water stress for the 12 plant descriptors.

| Parents | Number of leaves | Leaf area | Specific leaf weight | Leaf area index | Leaf water potential | Relative leaf water content | Photosynthetic efficiency | Proline content | Biological yield | Harvest index | Seed yield | Oil content |
| --- | --- | --- | --- | --- | --- | --- | --- | --- | --- | --- | --- | --- |
| CMS-XA | -4.43 | -0.11 | -0.06 | -0.61 | 0.001 | -0.75 | 0.79** | 0.10** | 8.76* | 0.65* | -1.3 | 1.40** |
| CMS-E002-91A | -2.35 | -0.12 | -0.19 | -0.67 | 0.07** | 5.09** | -0.46 | 0.07** | 26.47** | -1.3 | 1.75** | -1.45 |
| CMS-PKU-2A | -1.47 | 0.13** | 0.37** | 0.73** | -0.1 | 2.44** | -1.3 | -0.07 | -23.55 | 0.14 | 0.16 | -0.21 |
| CMS-ARG-2A | -3.45 | -0.02 | 0.09** | -0.11 | -0.11 | 2.06** | -2.01 | 0.02* | -40.94 | -1.42 | -2.90 | 1.29** |
| CMS-ARG-3A | -3.94 | -0.12 | -0.03 | -0.64 | 0.25** | 6.94** | -5.98 | -0.03 | 52.69** | 1.11** | 5.17** | -0.24 |
| CMS-ARG-6A | 2.33** | -0.03 | -0.12 | -0.31 | -0.08 | 1.50** | 0.77** | 0.03** | -30.93 | 2.65** | 7.21** | -0.37 |
| CMS-DV-10A | 0.12 | -0.03 | -0.04 | -0.12 | -0.04 | -3.09 | -3.09 | -0.08 | 8.97* | -0.38 | 0.21 | -0.75 |
| CMS-PHIR-27A | -0.34 | -0.13 | -0.28 | -0.7 | 0.21** | 5.03** | 1.62** | -0.02 | -5.51 | 1.91** | 1.00 | -0.81 |
| CMS-PRUN-29A | -0.41 | 0.16** | 0.32** | 0.89** | -0.22 | -5.61 | 3.57** | 0.08** | 48.11** | 0.33 | 1.07 | 1.94** |
| CMS-40A | 2.82** | -0.08 | -0.24 | -0.4 | 0.04* | -0.87 | 0.64** | -0.02 | -21.66 | 1.00** | 3.84** | -0.25 |
| CMS-42A | 1.86** | 0.14** | 0.18** | 0.82** | 0.14** | -8.02 | -2.12 | -0.06 | -9.99 | -0.52 | 3.18** | 0.66** |
| CMS-234A | 1.58** | 0.16** | -0.01 | 0.88** | 0.20** | -8.97 | 5.56** | 0.01 | 111.49** | -2.2 | 3.39** | 0.49** |
| CMS-38A | 1.29** | 0.09** | 0.07** | 0.43** | -0.18 | -7.76 | 1.25** | -0.01 | -81.18 | 0.05 | -7.13 | -0.65 |
| NC-41B (C) | -1.47 | -0.04 | -0.01 | -0.22 | -0.1 | 5.01** | 1.35** | -0.02 | 5.95 | 1.04** | -5.24 | -1.85 |
| 42B | 0.86** | 0.001 | -0.06 | 0.01 | -0.07 | 6.98** | -0.6 | 0.02* | -48.68 | -3.05 | -10.4 | 0.81** |
| RCR-8297 | 0.26** | -0.02 | -0.03 | -0.06 | 0.05** | -0.19 | 0.93** | 0.001 | -18.32 | 2.54** | 3.15** | -0.27 |
| P69R | 1.09** | 0.03** | 0.01 | 0.16** | -0.14 | 2.21** | 1.67** | 0.04** | 23.20** | -0.85 | -3.5 | -0.14 |
| P124R | -0.29 | -0.05 | -0.09 | -0.28 | 0.02* | -2.33 | -1.95 | -0.02 | -27.59 | 0.17 | -2.1 | -0.28 |
| P100R | -1.06 | 0.04** | 0.11** | 0.19** | 0.07** | 0.30** | -0.64 | -0.02 | 22.71** | -1.86 | 2.44** | 0.68** |
| *GCA Under Water Stress* |  |  |  |  |  |  |  |  |  |  |  |  |
| CMS-XA | -1.36 | -0.08 | -0.12 | -0.37 | -0.1 | 6.28** | -0.75 | 0.32** | 15.09** | 2.34** | 6.60** | 0.22** |
| CMS-E002-91A | 0.19 | 0.04** | 0.08** | 0.32** | 0.15** | -7.82 | -0.07 | 0.24** | -10.51* | 1.93** | 1.82** | 0.36** |
| CMS-PKU-2A | -0.98 | 0.10** | 0.27** | 0.64** | 0.12** | 4.08** | -1.42 | -0.22 | -49.58 | 6.01** | -0.8 | 1.86** |
| CMS-ARG-2A | -2.78 | -0.07 | -0.02 | -0.29 | 0.41** | 5.18** | 0.19 | 0.08** | -57.42 | 1.54** | -3.82 | 1.58** |
| CMS-ARG-3A | -1.79 | -0.17 | -0.31 | -0.85 | -0.14 | -2.72 | -1.79 | -0.06 | -13.07 | 1.59** | 4.80** | 3.23** |
| CMS-ARG-6A | 1.99** | 0.03** | 0.001 | -0.05 | -0.22 | -4.9 | 2.18** | -0.09 | 14.80** | 0.41 | 3.78** | -0.45 |
| CMS-DV-10A | -3.00 | -0.06 | 0.03 | -0.25 | -0.26 | -7.56 | -2.33 | -0.22 | -58.69 | 3.27** | -0.04 | 0.68** |
| CMS-PHIR-27A | -2.94 | -0.13 | -0.2 | -0.63 | -0.22 | 2.62** | -2.27 | -0.04 | -45.97 | 1.88** | -0.79 | -1.22 |
| CMS-PRUN-29A | -2.12 | 0.06** | 0.22** | 0.41** | -0.03 | -0.61 | -0.09 | 0.28** | 31.44** | 3.43** | 8.98** | 1.15** |
| CMS-40A | 3.92** | -0.02 | -0.11 | -0.19 | -0.16 | -4.99 | 0.80** | -0.19 | 74.96** | -2.94 | 0.69 | -0.52 |
| CMS-42A | 1.63** | 0.18** | 0.29** | 1.05** | -0.18 | 2.42** | -0.82 | -0.18 | -11.28 | -1.72 | 0.53 | -0.58 |
| CMS-234A | 1.53** | 0.09** | -0.1 | 0.59** | 0.20** | -2.04 | 4.68** | 0.03** | 93.32** | -2.84 | 3.44** | 0.27** |
| CMS-38A | 1.42** | 0.02* | -0.13 | -0.23 | 0.22** | -2.17 | 0.96** | 0.02* | -61.22 | -1.59 | -7.12 | -0.01 |
| NC-41B (C) | -4.81 | -0.06 | 0.15** | -0.27 | 0.21** | 11.77** | 1.50** | -0.03 | 7.63 | -7.67 | -15.49 | -4.71 |
| 42B | 3.10** | 0.08** | -0.07 | 0.11** | 0.01 | 0.47 | -0.78 | 0.06** | 70.51** | -5.63 | -2.59 | -1.85 |
| RCR-8297 | -0.27 | 0.001 | 0.04** | 0.08** | 0.01 | 1.58** | 1.34** | 0.03** | -28.1 | 2.88** | 1.70** | 0.34** |
| P69R | 0.96** | -0.01 | -0.06 | -0.03 | -0.05 | -2.96 | 0.75** | 0.07** | 60.37** | -3.15 | -1.05 | 0.32** |
| P124R | -0.13 | -0.04 | -0.08 | -0.26** | 0.16** | 0.50** | -1.46 | -0.03 | -21.67 | 1.38** | -1.06 | -0.35 |
| P100R | -0.56 | 0.05** | 0.10** | 0.21** | -0.11 | 0.88** | -0.64 | -0.07 | -10.6 | -1.11 | 0.41 | -0.31 |

#### 2.2.1. Number of leaves per plant

In case, of number of leaves the female parent CMS-ARG-6A recorded highly significant GCA effects (2.33 and 1.99) under both the environments and it belongs to wild species *H. argophyllus* (Table 3). Whereas, among the male parents (testers) RCR-8297 was recorded as a best general combiner (0.26) under the normal environment, while P69R (1.09 and 0.96) under both the environments (Table 3). Among sixty hybrids, twenty-five hybrids recorded significant positive SCA effects under normal environment; whereas, under stress environment, twenty-one hybrids recorded high and positive SCA effects (Table S1). The top three common hybrids combinations which recorded highly significant SCA effects under both the environments are CMS-38A × RCR-8297, CMS-ARG-6A × P124R, and CMS-40A × P124R (Table 4).

**Table 4.** Common three significant cross combinations with high specific combining ability (SCA).

| Traits | Crosses | N | S |
| --- | --- | --- | --- |
| Number of leaves | CMS-38A × RCR-8297 | 4.65** | 1.79** |
|  | CMS-ARG-6A × P124R | 3.08** | 2.65** |
|  | CMS-40A × P124R | 2.22** | 1.01** |
| Leaf area | CMS-ARG-2A × RCR-8297 | 0.42** | 0.38** |
|  | CMS-234A × P100R | 0.34** | 0.31** |
|  | CMS-XA × P69R | 0.23** | 0.03** |
| Specific leaf weight | CMS-ARG-2A × RCR-8297 | 0.88** | 0.82** |
|  | CMS-XA × P69R | 0.67** | 0.13** |
|  | CMS-234A × P100R | 0.44** | 0.45** |
| Leaf area index | CMS-ARG-2A × RCR-8297 | 2.25** | 2.01** |
|  | CMS-234A × P100R | 1.90** | 1.79** |
|  | CMS-38A × RCR-8297 | 1.09** | 1.13** |
| Leaf water potential | CMS-PRUN-29A × RCR-8297 | 0.45** | 0.13** |
|  | CMS-38A × RCR-8297 | 0.40** | 0.28** |
|  | 42B × P69R | 0.33** | 0.33** |
| Relative leaf water content | CMS-234A × RCR-8297 | 5.94** | 17.21** |
|  | CMS-E002-91A × P69R | 5.77** | 7.54** |
|  | CMS-40A × P69R | 5.70** | 7.52** |
| Photosynthetic efficiency | CMS-PHIR-27A × P100R | 4.21** | 1.18** |
|  | CMS-38A × RCR-8297 | 3.95** | 7.14** |
|  | CMS-E002-91A × P100R | 1.97** | 2.85** |
| Proline content | 42B × RCR-8297 | 0.20** | 0.62** |
|  | CMS-XA × P69R | 0.18** | 0.60** |
|  | CMS-PRUN-29A × P124R | 0.14** | 0.38** |
| Biological yield | CMS-38A × RCR-8297 | 98.05** | 58.42** |
|  | CMS-XA × P124R | 84.12** | 58.67** |
|  | CMS-ARG-6A × P124R | 67.40** | 184.79** |
| Harvest index | CMS-PHIR-27A × RCR-8297 | 9.10** | 2.47** |
|  | 42B × P100R | 5.39** | 2.83** |
|  | CMS-ARG-3A × P124R | 5.18** | 4.42** |
| Seed yield | CMS-ARG-2A × P100R | 9.97** | 10.70** |
|  | CMS-PKU-2A × P124R | 8.31** | 7.46** |
|  | 42B × P69R | 8.14** | 11.60** |
| Oil content | CMS-E002-91A × P124R | 2.11** | 1.62** |
|  | NC-41B (C) × RCR-8297 | 2.08** | 3.32** |
|  | CMS-PRUN-29A × P100R | 2.03** | 1.77** |

\*\*, \* indicate significant at $p < 0.01$ , $p < 0.05$ , respectively.

#### 2.2.2. Leaf area (m^2^)

Two CMS analogues CMS-PKU-2A (*H. annuus*) and CMS-PRUN-29A (*H. praecox* ssp. Runyonii) recorded a highly significant GCA effect under both the environments for leaf area (Table 3). Among, the testers P69R was recorded as a significant combiner under normal environment while P100R was recorded significant under both the environments (Table 3). Among hybrids, twenty-three hybrids recorded significant positive SCA effect both the environments (Table S1). The top three common hybrids for SCA both of the environments were CMS-ARG-2A × RCR-8297, CMS-234A × P100R and CMS-XA × P69R (Table 4).

#### 2.2.3. Specific leaf weight (g)

In case of specific leaf weight, CMS-PKU-2A recorded highly significant GCA under normal as well as in water stress environment (Table 3). The male parent P100R was observed as a good combiner for this trait under both the environments. A total of eighteen hybrids recorded highly significant positive SCA effects under both the environments (Table S1). Whereas, the top three common combinations for SCA were CMS-ARG-2A × RCR-8297, CMS-XA × P69R, and CMS-234A × P100R (Table 4).

#### 2.2.4. Leaf area index

For leaf area index the female parent CMS-PRUN-29A (0.89) from wild sources and CMS-42A, CMS-234A and CMS-38A from PET-1 source were recorded as significant general combiners (Table 3). Among male parents, P100R was observed with a significant GCA effect (0.19 and 0.21) under both environments (Table 3). Out of 60 hybrids, nineteen were recorded highly significant SCA for LAI under both the environments (Table S1), although top three common combinations for SCA were CMS-ARG-2A × RCR-8297, CMS-234A × 100R and CMS-38A × RCR-8297 (Table 4).

#### 2.2.5. Leaf water potential (MPa)

For leaf water potential under the normal environment, CMS-ARG-3A was observed as significant combiner (Table 3). Under stress environment, CMS-E002-91A, CMS-PKU-2A and CMS-ARG-2A, CMS-234A, CMS-38A and NC-41B were observed as better general combiners for leaf water potential (Table 3). A total of twenty-one hybrids were reported to have significant positive SCA effects under normal environment whereas, twenty-seven hybrids had the good combining ability under water stress environment (Table S1). The best three cross combinations identified with high SCA effects for leaf water potential under both the environments were CMS-PRUN-29A × RCR-8297, CMS-38A × RCR-8297, and 42B × P69R respectively (Table 4).

#### 2.2.6. Relative leaf water content (%)

The six CMS analogues out of nine and both the maintainers were recorded as significant general combiners for relative leaf water content were identified as good combiners under the normal environment (Table 3). In water stress environment, female lines CMA-XA, CMS-PKU-2A, CMS-ARG-2A and CMS-PHIR-27A from wild sources and CMS-NC-41B from the PET-1 source were observed as significant combiners (Table 3). A total of twenty-eight hybrids recorded significant positive SCA effects in the normal environment. Twenty-nine hybrids recorded significant positive SCA effects under stress environment (Table S1). The three cross combinations viz. CMS-234A × RCR-8297, CMS-E002-91A × P69R, and CMS-40A × P69R were identified with highly significant SCA effects for relative leaf water content under both the environments (Table 4).

#### 2.2.7. Photosynthetic efficiency (SPAD reading)

In the normal environment and water stress environment, the female parents CMS-234A was observed with the highest GCA effect (5.56 and 4.68 1) (Table 3). While the male parent RCR-8297 and P69R were recorded as a significant combiners (0.93 and 1.34; 1.67 and 0.75) under both the environments for photosynthetic efficiency (Table 3). Twenty-one cross combinations recorded significant positive SCA effects in a normal environment. In the water stress condition, seventeen cross combinations were identified with high SCA values (Table S1). The cross combinations viz. CMS-PHIR-27A × P100R, CMS-38A × RCR-8297 and CMS-E002-91A × P100R were identified with high SCA effects for photosynthetic efficiency under both the environments (Table 4).

#### 2.2.8. Proline content (mg/g of dry weight)

The CMS analogues CMS-XA, CMS-E002-91A, CMS-ARG-6A and CMS-PRUN-29A were recorded very good combiners for the proline content in a normal environment (Table 3). Whereas, under the stress environment, CMS-XA, CMS-E002-91A, CMS-ARG-3A and CMS-PRUN-29A and CMS-234A were observed as a good general combiner (Table 3). The tester P69R appeared to be the significant general combiner for proline content under both the environments. Best three hybrids 42B × RCR-8297, CMS-XA × P69R and CMS-PRUN-29A × P124R were recorded highly significant positive SCA effects under both the environments (Table 4).

#### 2.2.9. Seed yield (g/plant)

For the seed yield under normal environment the CMS analogues CMS-E002-91A (1.75), CMS-ARG-3A (5.17), CMS-ARG-6A (7.21) and CMS-40A (3.84) from wild sources; whereas, CMS-42A (3.18) and CMS-234A (3.39) (from PET-1) were recorded as significant for the GCA effects (Table 3). Whereas, under water-limited conditions, the CMS analogues CMS-XA (6.60), CMS-E002-91A (1.82), CMS-ARG-3A (4.80), CMS-ARG-6A (3.78), CMS-PRUN-29A (8.98) and CMS-234A (3.44) had highly significant GCA effects for seed yield (Table 3). The CMS analogues CMS-E002-91A, CMS-ARG-2A, CMS-ARG-3A and CMS-234A recorded for significant combining ability under both the environments. Twenty hybrids recorded highly significant positive SCA effects under normal environment while twenty-six hybrids reported positive SCA effects under stress environment (Table S1).

#### 2.2.10. Biological yield (g/plant)

Lines CMS-DV-10A and CMS-XA were good combiner for biological yield from the wild sources under a normal environment (Table 3). Whereas, under water stress environment, the female lines CMS-XA, CMS-ARG-6A and CMS-PRUN-29A were recorded as significant combiners while CMS-E002-91A observed as a significant combiner for this trait. CMS-234A (111.49 and 93.32) and CMS-PRUN-29A (48.11 and 31.44) were recorded as significant for biological yield per plant under both normal and stress environments (Table 3). The tester P69R appeared to best among others for biological yield per plant under both the environments (23.20 and 60.37). Best three common hybrids combinations for SCA were CMS-38A × RCR-8297, CMS-XA × P124R and CMS-ARG-6A × P124R respectively (Table 4).

#### 2.2.11. Harvest index (%)

All CMS analogues from wild sources were recorded as very good combiner except CMS-ARG-6A whereas, from conventional PET-1 source, none of the female lines had significant GCA for harvest index under water stress environment (Table 3). The tester RCR-8297 appeared to be a good general combiner for harvest index with significant and positive GCA effects (2.54 and 2.88) under both the environments (Table 3). Twenty-four hybrids under normal environment and twenty hybrids under water stress environment recorded significant positive SCA effects (Table S1). Best three hybrids were CMS-PHIR-27A × RCR-8297, 42B × P100R and CMS-ARG-3A × P124R for significant SCA effects for harvest index common to both environments (Table 4).

#### 2.2.12. Oil content (%)

In case of oil content, the analysis pooled over the years under different water regimes showed that CMS analogues CMS-XA (1.40), CMS-ARG-2A (1.29), CMS-PRUN-29A (0.75), CMS-42A (0.66) and CMS-234A (0.49) were recorded as significant general combiners under the normal environment (Table 3). Whereas, in water stress environment, all CMS analogues except CMS-ARG-6A and CMS-PHIR-27A were observed significant combiner for oil content (Table 3). The male parent RCR-8297 and P69R were recorded as highly significant general combiner under stress environment, while P100R was recorded as a significant general combiner under the normal environment (Table 3). Twenty-two hybrids under normal environment and twenty-eight hybrids under stress environment recorded significant positive SCA effects (Table S1).

### 2.3. Correlations

In case of normal environment, 16 correlation coefficients were found to be significant (P < 0.05) (Figure 2). Among them four were negative correlations and three were absolute correlations. Harvest index was found negatively correlated with biological yield, specific leaf weight, leaf area leaf area index (Figure 2). While absolute correlations exist between leaf area and leaf area index. Also, specific leaf weight with leaf area and leaf area index (0.92) (Figure 2). Whereas, under water stress environment, a total of 14 correlation coefficients were found to be significant (P < 0.05) (Figure 4). Thirteen were positive correlations and one was a negative correlation i.e. between harvest index and biological yield (−0.59) (Figure 3). Also, the specific leaf weight was absolutely correlated with leaf area (0.90) and leaf area index (0.92) (Figure 3). The seed yield was found correlated with the oil content (0.49), number of leaves (0.44), and biological yield (0.43) (Figure 4). Number of leaves and biological yield (0.46) was found correlated. Leaf water potential was correlated with leaf area, leaf area index and specific leaf weight in water-limited conditions (Figure 3).

**Figure 2.**
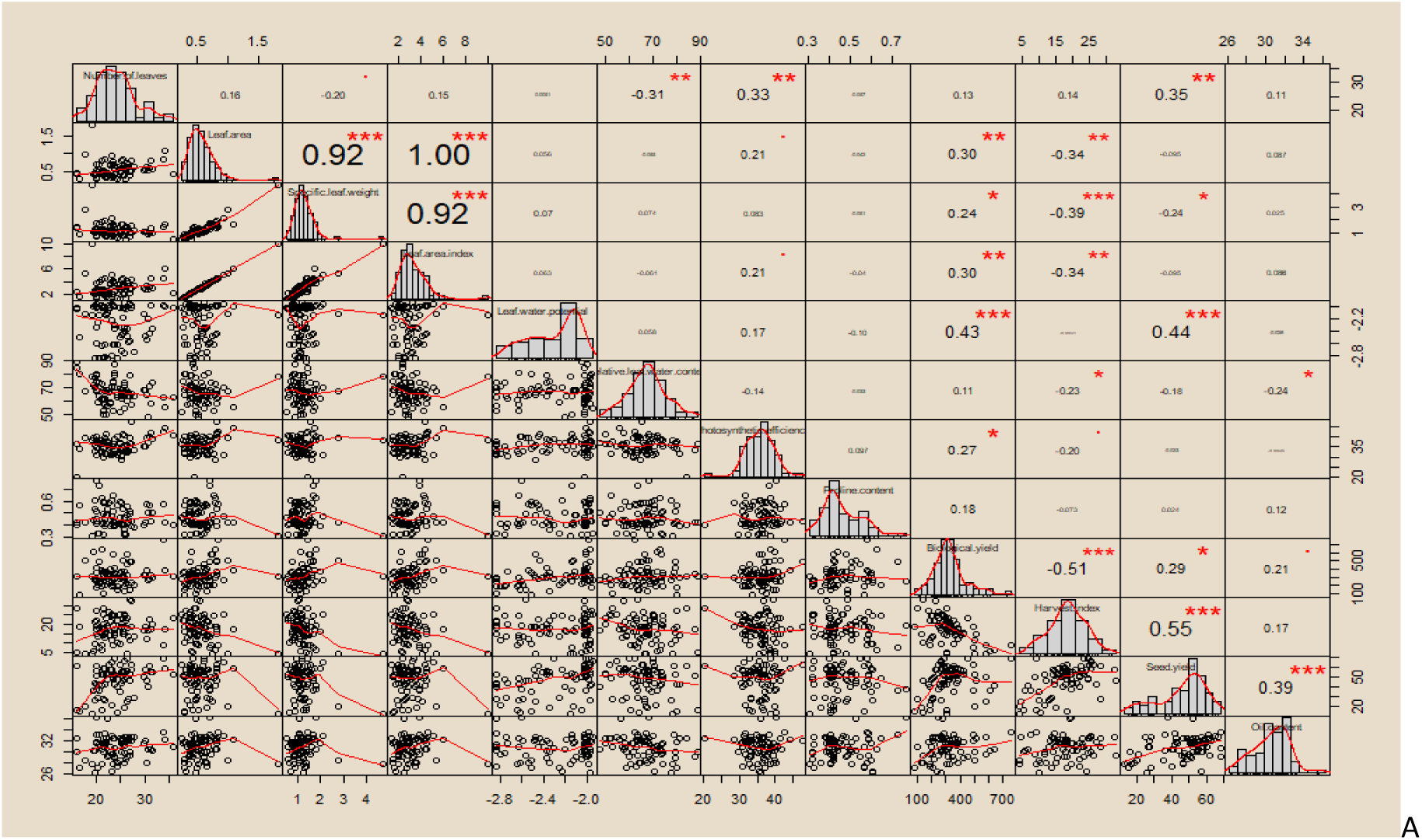
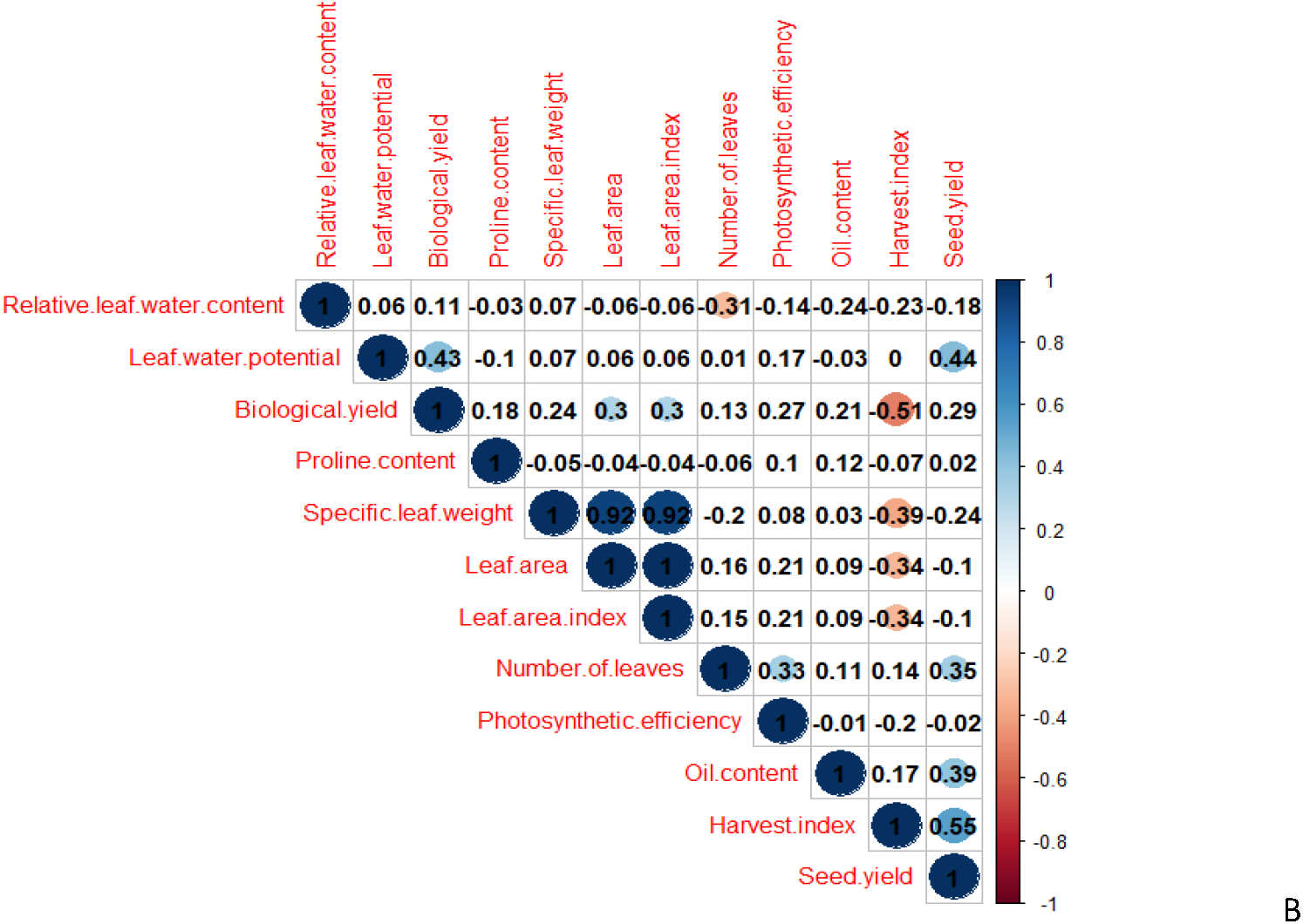
Under normal condition Pearson’s correlation coefficients with significant values at p<0.001 (***), p<0.01 (**), or p<0.05 (*), respectively (upper diagonal) along with the pattern of the distribution of data via scattering plots and histograms (lower diagonal) (A). While all the significant Pearson’s correlation coefficients at p<0.05 (B).

**Figure 3.**
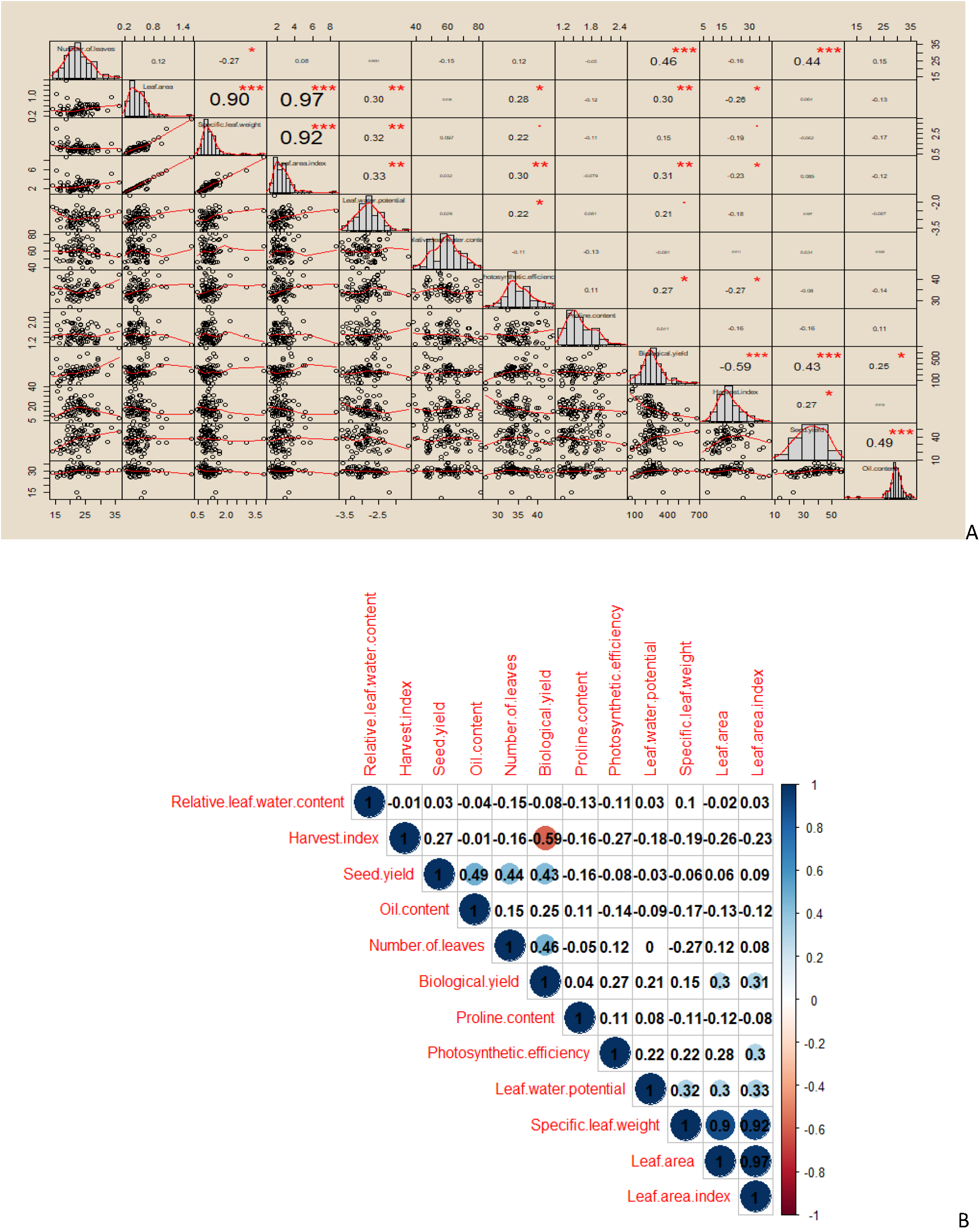
Under water stress Pearson’s correlation coefficients with significant values at p<0.001 (***), p<0.01 (**), or p<0.05 (*), respectively (upper diagonal) along with the pattern of the distribution of data via scattering plots and histograms (lower diagonal) (A). While all the significant Pearson’s correlation coefficients at p<0.05 (B).

**Figure 4.**
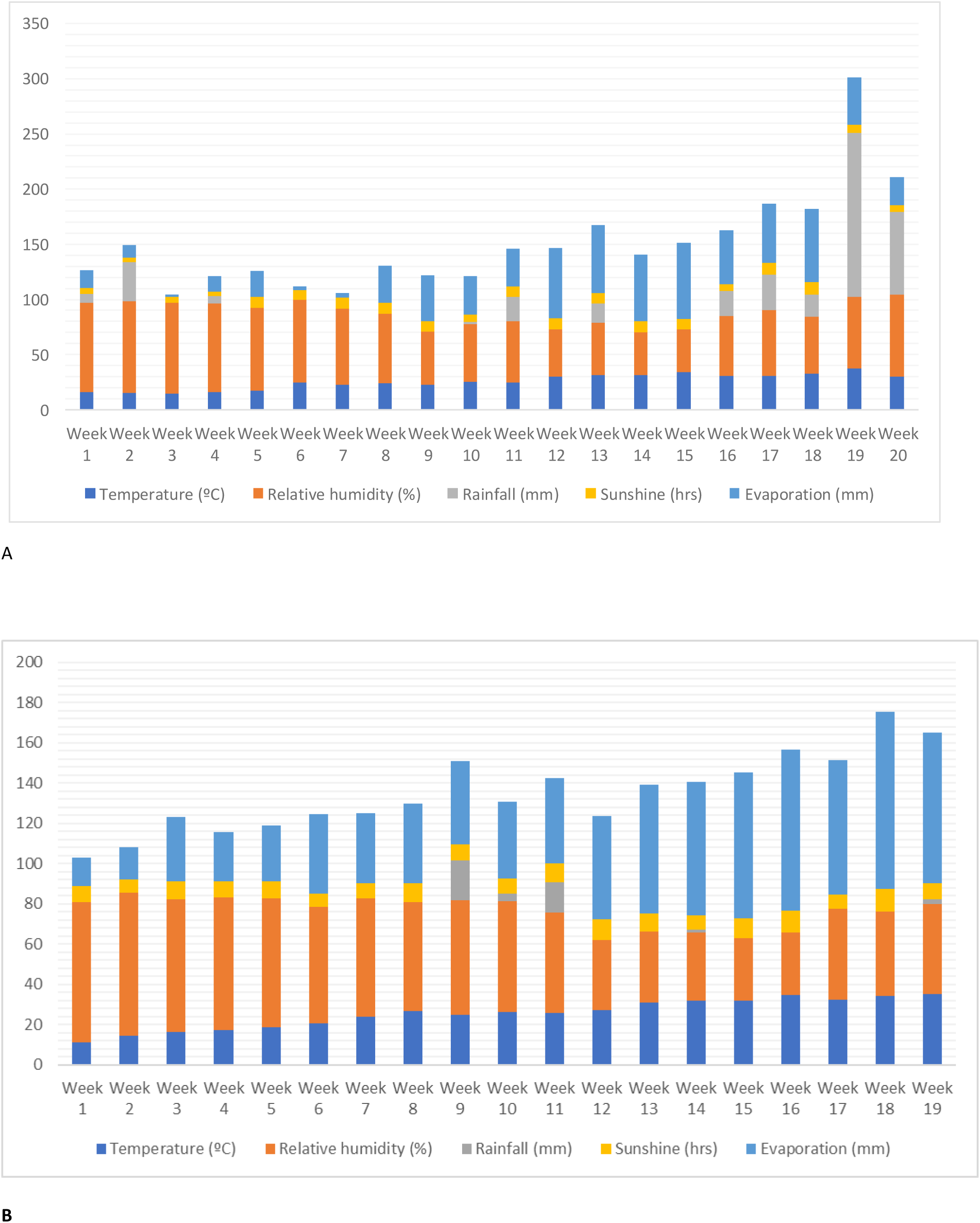
Weather parameters during first (A) and second (B) year.

## 3. Discussion

Wild species are recognized as important sources of valuable genes for the breeding objective of drought tolerance [34–36]. In major crops (cereals) the important genes for drought tolerance are well identified, and further, they are introgressed into the genome of popular cultivated varieties[37,38]. Whereas, in sunflower, only H. argophyllus has been extensively studied than any other species of this genus for drought tolerance [23,39]. Most of the times when a breeding experiment was performed under normal conditions, the wild relatives used were responsible for the lower improvement of yield and quality [40]. Whereas, under drought environment, the yield gain and performance improvement due to the use of wild relatives is more prominent. In the case of sunflower, the problem of homogeny of cytoplasm could lead to potential problems in the future [41]. Also, the balanced effect of cytoplasm and nuclear genome need to be well studied before commercialization of any sunflower variety as sometime such interactions could result in lower yields [42].

In our study the 13 CMS lines, 2 maintainer lines along with 4 restorer lines, and 60 F1 hybrids along with interspecific hybrids of six wild species. This study is among the largest studies carried out in sunflower to study cytoplasmic effects along with drought tolerance. Mostly, we have identified cross combinations with wild species are more significant under water-limited situations. In case of drought, we have found that proline content was three times more in the leaves than compared to the normal conditions. These results are quite similar to what people have found in sunflower and in other oilseeds [24,43].

The overall significant differences in mean values showed a significant amount of variation for all the studied traits. It was clearly observed that different cytoplasmic sources significantly influenced traits under both conditions. Earlier studies pointed out the effect of cytoplasm sources on different traits and it was concluded that the cytoplasmic genome affects qualitative and quantitative traits [44–46]. SCA effects are more important for cross-pollinated plants, whereas CGA effects are more important for self-pollinated plants [47]. Being a cross-pollinated plant sunflower showed significantly higher SCA values than corresponding GCA values for most of the 12 descriptors. Moreover, the importance of combining ability in a selection of parents for hybridization has been well emphasized by many workers in sunflower [48,49]. The good general combiner females lines crossed with good general combiner (testers) to yield hybrids, synthetics and composite varieties[50].

The non-additive gene effects were identified governing the inheritance of important traits like leaf area, specific leaf weight, leaf area index, leaf water potential, relative leaf water content, photosynthetic efficiency, proline content, seed yield, harvest index and oil content under normal and in water stress environment. In this direction, earlier studies have reported non-additive gene action for seed yield and oil content while additive gene action for oil content [51]. In contrast, some studies also reported additive gene action for the inheritance of seed yield, oil content etc.[52]. Overall, The female parents (lines) had a large proportion in the expression of traits than male parents also affirms the role of cytoplasmic and nuclear interactions in the expression of these traits. This also shows the importance of plant material diversity used for similar studies [53]. Similarly, significant GCA effect for seed yield was stated in the previous studies in that parent which were good general combiners for economic traits might be extensively used in hybridization programs to identify suitable parents for hybridization and developing potential hybrids for seed yield [54,55]. The female lines CMS-234A from cultivated source and CMS-XA, ARG-2A and PRUN-29A from the wild source appeared as good combiners for oil content. Among testers RCR-8297 for seed yield under both the environments while RCR-8297 and P69R for oil content under stress environment with high positive GCA estimates are desirable parents to be used for developing sunflower hybrids with improved oil content [52, 56,57].

The hybrids CMS-XA x P100R, CMS-ARG-2A x P100R, and 42B x RCR-8297 were considered as the best specific combiners for seed yield and CMS-E002-91A x P124R, CMS-E002-91A x P100R, and CMS-40A x P124R for oil content. Higher positive SCA effects indicated that these characters are influenced by the dominance and overdominance gene actions. In consonance to our findings, higher positive SCA for head diameter; for 1,000-achene weight; and for seed yield and yield-related traits and for oil content was previously reported [58-60] The heterotic performance of hybrid combinations depends upon the combining abilities of their parents [59]. The superior hybrids were obtained by crossing CMS females and restorer males with high GCA and SCA effects [54]. The overall variation among CMS lines for studied traits were greater than the restorer lines indicating some degrees of maternal effect on traits, particularly for seed and oil yield [61]. The seed yield was found correlated under both the environments to harvest index, number of leaves and oil content. While, proline content was not correlated with any of the descriptors. Whereas, under biotic stress, the yield was independent of other traits[60].

## 4. Materials and Methods

### 4.1. Experimental Layout and Material

The open experimental fields were settled at the research farm of Punjab Agricultural University, Ludhiana, India (coordinates at 30°54’6″ N 75<u>°</u>;48’27″ E). The experimental material comprised 19 genotypes of sunflower in the form of different species (Table 5). Among them, 9 were alloplasmic CMS (A) lines, 4 euplasmic CMS lines (A), 2 maintainer lines (B), and 4 restorer lines (R) (Table 5). Both, A lines (13) and B lines (2) were crossed with the R lines (4) to produce sixty one-way F1 hybrids (Table 5). The hybrids were produced as the nine alloplasmic lines and euplasmic CMS lines from petiolaris source were crossed with restorers to synthesize a set of 52 A × R crosses (TABLE 5). Subsequently, both maintainer lines were made sterile using gibberellic acid (GA_3_) (100 ppm) at the star bud initiation stage for three consecutive days [62, 63]. Thereafter, these maintainer lines were also crossed with the restorer lines to synthesize a set of eight B × R crosses (Table 1). The study was conducted over the period of two years i.e. 1^st^ year during February 2011 and 2nd year in the February 2012. In the first year of the experiment, a total of 52 A × R crosses and 8 B × R crosses along with parental lines were planted in February 2011 via a randomized complete block design (RCBD) with three replications. In the field, each parental genotype (19) and cross combination (60) was represented by a plot of two rows (3 m^2^ each) with an inter and intra-row spacing maintained at 60 cm and 30 cm, respectively. There were 20 plants (10 in each plot) of every parent and their hybrids in each replication. Similarly, the nearby field was chosen for the drought screening trial, the same set of experiment was repeated with a different water regime by withholding irrigation during anthesis and soft dough stages of crop growth [64]. The same experiment was repeated in the February 2012 for the second-year evaluation trial.

**Table 5.** List of sunflower accessions along with their source (species) and hybrids used for the characterization.

| A/B/R lines accessions | Species | Hybrids |  |  |  |
| --- | --- | --- | --- | --- | --- |
|  |  | RCR-8297 | P69R | P124R | P100R |
| A Lines (Alloplasmic) |  |  |  |  |  |
| CMS-XA | Unknown | CMS-XA × RCR-8297 | CMS-XA × P69R | CMS-XA × P124R | CMS-XA × P100R |
| CMS-E002-91A | H. annuus | CMS-E002-91A × RCR-8297 | CMS-E002-91A × P69R | CMS-E002-91A × P124R | CMS-E002-91A × P100R |
| CMS-PKU-2A | H. annuus | CMS-PKU-2A × RCR-8297 | CMS-PKU-2A × P69R | CMS-PKU-2A × P124R | CMS-PKU-2A × P100R |
| CMS-ARG-2A | H. argophyllus | ARG-2A × RCR-8297 | CMS-ARG-2A × P69R | CMS-ARG-2A × P124R | CMS-ARG-2A × P100R |
| CMS-ARG-3A | H. argophyllus | CMS-ARG-3A × RCR-8297 | CMS-ARG-3A × P69R | CMS-ARG-3A × P124R | CMS-ARG-3A × P100R |
| CMS-ARG-6A | H. argophyllus | CMS-ARG-6A × RCR-8297 | CMS-ARG-6A × P69R | CMS-ARG-6A × P124R | CMS-ARG-6A × P100R |
| **CMS-DV-10A** | *H. debilis ssp. Vestitus* | CMS-DV-10A × RCR-8297 | CMS-DV-10A × P69R | CMS-DV-10A × P124R | CMS-DV-10A × P100R |
| **CMS-PHIR-27A** | *H. praecox ssp. Hirtus* | CMS-PHIR-27A × RCR-8297 | CMS-PHIR-27A × P69R | CMS-PHIR-27A × P124R | CMS-PHIR-27A × P100R |
| **CMS-PRUN-29A** | *H. praecox ssp. Runyonii* | CMS-PRUN-29A × RCR-8297 | CMS-PRUN-29A ×<br>P69R | CMS-PRUN-29A ×<br>P124R | CMS-PRUN-29A × P100R |
| **A Lines (Euplasmic)** |  |  |  |  |  |
| **CMS-40A** | *H. petiolaris (conventional)* | CMS-40A × RCR-8297 | CMS-40A × P69R | CMS-40A × P124R | CMS-40A × P100R |
| **CMS-42A** | *H. petiolaris (conventional)* | CMS-42A × RCR-8297 | CMS-42A × P69R | CMS-42A × P124R | CMS-42A × P100R |
| **CMS-234A** | *H. petiolaris (conventional)* | CMS-234A × RCR-8297 | CMS-234A × P69R | CMS-234A × P124R | CMS-234A × P100R |
| **CMS-38A** | *H. petiolaris (conventional)* | CMS-38A × RCR-8297 | CMS-38A × P69R | CMS-38A × P124R | CMS-38A × P100R |
| **B Lines (Maintainer)** |  |  |  |  |  |
| **NC-41B** | *H. petiolaris (conventional)* | NC-41B × RCR-8297 | NC-41B × P69R | NC-41B × P124R | NC-41B × P100R |
| **42B** | *H. petiolaris (conventional)* | 42B × RCR-8297 | 42B × P69R | 42B × P124R | 42B × P100R |
| **R Lines (Restorer)** |  |  |  |  |  |
| **RCR-8297** | *H. annuus* |  |  |  |  |
| **P69R** | *H. annuus* |  |  |  |  |
| **P124R** | *H. annuus* |  |  |  |  |
| **P100R** | *H. annuus* |  |  |  |  |

### 4.2. Weather Parameters and Soil Properties

The meteorological data (weekly intervals) during the entire crop season of 2011 (February 2011 to June 2011) and 2012 (February 2012 to June 2012) is represented in Figure 4. The air temperature ranged from 4.2°C to 43.7°C (Figure 4).

Physical properties of soil were also analyzed to determine the native fertility and soil texture of the experimental fields (Table 6). The composite soil samples were collected from the topsoil layer (0–15 cm) of the experimental site and analyzed for sand, silt clay content and soil temperature using the methods defined elsewhere [65].

**Table 6.**
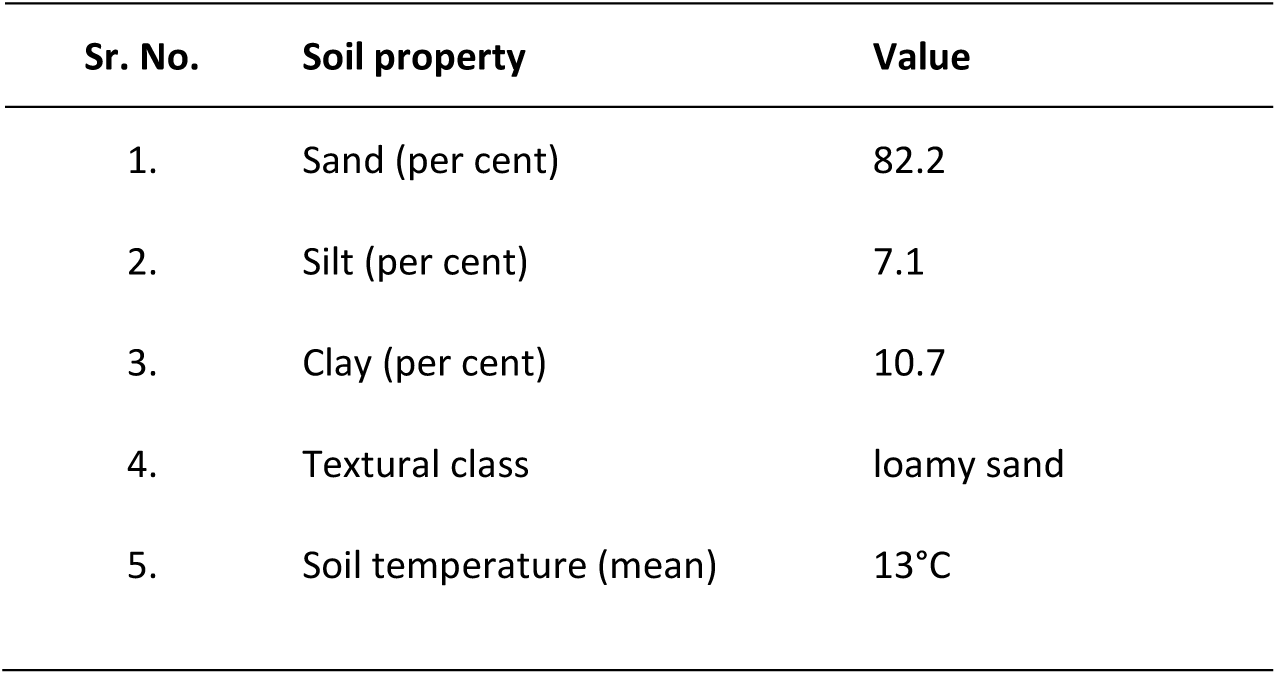
Soil physical properties of experimental site.

| Sr. No. | Soil property | Value |
| --- | --- | --- |
| 1. | Sand (per cent) | 82.2 |
| 2. | Silt (per cent) | 7.1 |
| 3. | Clay (per cent) | 10.7 |
| 4. | Textural class | loamy sand |
| 5. | Soil temperature (mean) | 13°C |

### 4.3. Characterization of Plants and Data Analysis

Five central plants per plot (i.e. ten plants per replication) were used for the data acquisition and characterization. The number of leaves per plant was determined as the average of plant leaves at the time of maturity. The leaf area (m^2^) was calculated using an area meter (LI-3100; LICOR, Lincoln, NE, USA).

Whereas, the specific leaf weight (g) was estimated as total leaf weight divided by the number of leaves. The leaf area index (LAI) was estimated using the formula, LAI= (A×N)/10,000, where A is leaf area (cm^2^) and N is plants per m^2^. [66]. The leaf water potential in Mega-Pascal’s (MPa) was recorded with the help of vapour pressure osmometer (Wescor Vapro 5520, Wescor Inc. Logan, UT, USA). The relative leaf water content (RLWC) was determined based on the formula, RLWC= 100 × (Fresh weight - Dry weight / Saturated weight - Dry weight) [67]. The chlorophyll content of leaves was recorded as a SPAD meter reading with the help of SPAD meter (SPAD 502 Plus Chlorophyll Meter, Minolta, Japan) [68]. The proline content (mg/g dry weight of leaf), was estimated based on the method defined elsewhere[69]. The biological yield (g/plant) was calculated as the above-ground weight of total plant including the sunflower head. Whereas, the harvest index (HI) was estimated based on the formula, HI= 100 × seed yield / total biomass [vegetative mass (VM) + Seed Yield] at maturity [70]. Further, to determine the seed yield the harvested seeds from five competitive plants were weighed and averaged. Oil content (%) was estimated using the nuclear magnetic resonance (NMR) analyzer (Newport Analyzer MK III A, Newport Instruments Ltd., Milton Keynes, England) [71].

The replicated data for all the descriptors across the period of two years was subjected to the statistical analysis. The popular (Unweighted Pair Group Method with Arithmetic Mean) UPGMA method of hierarchical clustering was used to visualize how parental genotypes are related based on the differences in the data of twelve descriptors using the R platform (R Core Team 2015). The “average” method was used with the function hclust [72]. The Line × Tester calculations were performed following standard procedures for the estimation of components of genetic variation, with the help of the SAS software, version 9.1.SAS Institute, Inc. SAS user’s guide. Version 9, 4th ed. Cary, NC. 2004 [32]. While the Pearson correlation coefficients were determined and plotted via packages corrplot and PerformanceAnalytics [73,74]. All the other remaining statistical analysis were carried out using the Statgraphics Centurion XVI software package (StatPoint Technologies, Warrenton, VA, USA).

## 5. Conclusions

The accession CMS-234A was the best general combiner for biological yield and photosynthetic efficiency under both the conditions. The cross combinations CMS-ARG-2A × RCR-8297, CMS-234A × P124R, and CMS-38A × P124R were found significant for biological yield, seed yield and oil content under both environments. Similarly, ARG-6A x P124R recorded maximum significant SCA effects for the number of leaves/plants. The hybrids CMS-ARG-2A x RCR-8297 and CMS-234A x P100R recorded highly significant SCA effects for leaf area, specific leaf weight and leaf area index under both the environments. For proline content two hybrids i.e. CMS-XA x P69R and CMS-ARG-2A x P124R shown the higher SCA effects under both the environments. The hybrids PRUN-29A x P69R and 234A x P69R recorded high SCA effects for biological yield/plant under both the environments. The parents (female and male) generally had high GCA (good combiners) for both the environments. Therefore, they can be exploited in the hybrid development program for drought tolerance, high yielding, and physiologically efficient with a diverse cytoplasmic background. All in all, the information about different cytoplasmic sources and their effect on traits under water-limited environment will be useful for the development of new hybrids adapted to the challenges imposed by the drought.

## Abbreviations

A: Cytoplasmic male sterile line
B: Maintainer line
CMS: Cytoplasmic male sterile
GA_3_: Gibberellic acid
GCA: General combining ability
HI: Harvest index
LAI: Leaf area index
R: Restorer line
RCBD: Randomized complete block design
RLWC: Relative leaf water content
SCA: Specific combining ability

